# TRIM 25 nuclear translocation is hampered by localization to influenza A virus non-structural protein 1 cytosolic aggregates

**DOI:** 10.1101/2022.10.10.511522

**Authors:** Anne Weiß, Ivan Haralampiev, Susann Kummer

## Abstract

Non-structural protein 1 (NS1) of influenza A virus (IAV) is involved in multiple blocking processes of cellular antiviral action. One of these includes the binding of cytosolic tripartite motif-containing protein 25 (TRIM25) to impede nuclear translocation and, thus, counteracting the block of viral RNA elongation taking place inside the nucleus. In addition, TRIM25 is part of the RIG-I signaling pathway and is an important component regarding the activation of interferon-induced genes and triggering cellular antiviral responses. In this study, we focus on the spatial localization of TRIM25 with respect to cytosolic NS1 aggregates found in IAV infected cells. Besides strain specific characteristics, we confirmed co-localization of TRIM25 to NS1 cytosolic aggregates at 16 hours post infection for all investigated IAV strains based on image analysis on single cell level. Immunoprecipitation further confirmed the interaction between NS1 and TRIM25 when comparing wt IAV and recombinant IAV devoid of NS1 expression. Of note, we combined fluorescence *in situ* hybridization targeting viral RNA with immunolabeling of TRIM25 suitable for nanoscopy and the diffraction limited detection of NS1 in one imaging approach. Our study reveals strain dependent differences in the block of IRF3 activation in respect to the accumulation of TRIM25 in cytosolic NS1 aggregates.

## 1. Introduction

The severity and diversity of the influenza A virus (IAV), which is the cause of infectious diseases of the upper and lower respiratory tract in humans, leads to seasonal influenza epidemics with peak values for each hemisphere per year (Molinari et al. 2007; Oxford 2000; Potter 2001). Modifications in virulence, acute risk from zoonotic reservoirs (e.g. birds, pigs) and/or the increase in antiviral resistance cause high economic costs and a significant burden on health systems worldwide (Fouchier et al. 2005; Gambotto et al. 2008; Ge et al. 2010; Salomon and Webster 2009). In order to address the problematic increase in antiviral drug resistance, recent infection research efforts have focused on combating viral ribonucleic acid (RNA) species. Imaging studies that clarify the question of whether viral RNA is orchestrated in the virus-induced block of cellular antiviral signal transmission provide valuable insights into complex RNA-protein interactions.

IAV strains are divided into subtypes according to two genes that form the most important surface proteins: haemagglutinin (HA) (Bouvier and Palese 2008; Skehel et al. 1982) and neuraminidase (NA) (Bouvier and Palese 2008; Chen et al. 2007; Nayak, Hui, and Barman 2004). Multiple copies of both proteins and the ion channel M2 (Holsinger and Lamb 1991; Pinto, Holsinger, and Lamb 1992) are embedded in the lipid bilayer of virions that display a spherical to filamentous shape (Harris et al. 2006). Underneath the envelope, matrix protein 1 forms a layer (Ruigrok et al. 2000; Ruigrok, Calder, and Wharton 1989) to protect the viral genome, which is composed of eight single-stranded RNAs in negative sense orientation (Bouvier and Palese 2008), complexed with the IAV nucleoprotein forms higher ordered structures: ribonucleoproteins (Baudin et al. 1994; Jennings et al. 1983).

Among the IAV proteins, the non-structural protein 1 (NS1) has a low abundance in the virion itself, but is essential for efficient viral replication (Bouvier and Palese 2008; Hutchinson et al. 2014). It is noteworthy that NS1 activity is strongly associated with reduced cellular antiviral defense: exemplarily, it (i) binds to dsRNA suppressing viral RNA degradation (Bornholdt and Prasad 2006; Li et al. 2004), (ii) inhibits the cellular transcription and the export of cellular transcripts from the nucleus (Fortes, Beloso, and Ortin 1994; Lu, Qian, and Krug 1994; Satterly et al. 2007), (iii) interacts with double-stranded RNA and/or RNA helicase, retinoic acid-inducible gene I (RIG-I) to block activation of various transcription factors, which initializes the expression of antiviral active factors (Guo et al. 2007; Mibayashi et al. 2007; Pichlmair et al. 2006) and (iv) amplifies this block by interaction with a tripartite motif containing protein 25 (TRIM25), an ubiquitin ligase which is a mediator of innate virus recognition by the interferon regulatory factor 3 (IRF3) pathway (Mibayashi et al. 2007). Once IRF3 was activated by phosphorylation, its localization shifts from the cytosol to the nucleus. Previous studies show that NS1 and TRIM25 bind and colocalize directly in cytosolic aggregates, preventing RIG-I activation, and downstream facilitation of interferon signalling (Mibayashi et al. 2007; Gack et al. 2009). Here, we investigated the complex NS1-TRIM25 interaction in cells with and without IRF3 activation for H1N1 and H3N2 IAV strains.

## 2. Materials and Methods

### Cells and viruses

Cell lines were maintained in Dulbecco’s modified Eagle Medium (DMEM) (Invitrogen) supplemented with 10 % foetal calf serum (Invitrogen), penicillin (200 U/ml)/streptomycin sulphate (100 μg/ml) (Capricorn scientific GmbH). Cells were maintained in a humidified incubator with 5 % CO_2_ at 37 C°. For maintaining stable transfected cell lines, 10 mg/ml puromycin was added to the culture medium. Cells subjected for immunofluorescence were seeded on cover slips in a 24 well plate format and for co-immunoprecipitation in 6 cm dishes with 80 % confluency on day of infection, respectively. Cells were infected with A/Hong Kong/1/1968 (HK/1/68, H3N2) (Stech et al. 2008), A/Panama/2007/1999 (Pan/2007/99, H3N2) (Matthaei, Budt, and Wolff 2013) and A/Puerto Rico/8/1934 (PR/8/34, H1N1, NCBI:txid211044) as well as corresponding ΔNS1 (Matthaei, Budt, and Wolff 2013; Garcia-Sastre et al. 1998) variants with a multiplicity of infection (MOI) of 1 for indicated times, respectively.

HK/1/68 and PR/8/34 were recovered in MDCK cells via reverse genetics as described elsewhere (Martinez-Sobrido and Garcia-Sastre 2010). If applicable, virus stocks were titrated by plaque assay as previously described (Morrill et al. 2010). In brief, serial 10-fold dilutions of virus stocks were prepared in DMEM supplemented with 0.3 % bovine serum albumin (BSA), 20 mM HEPES, penicillin (200 U/ml)/streptomycin sulphate (100 μg/ml) (Capricorn scientific GmbH) and TPCK-trypsin (2 μg/ml) (sigma-aldrich). MDCK or MDCKII NS1 (provided by Richard E. Randell, made simultaneously as described for A549 (Jackson et al. 2010)) cell monolayers in a 6 well plate format were incubated with 200 μl of each dilution for 1 h at 37°C, respectively. Subsequently cells were overlaid with 2 ml of 2.4 % Avicel (FMC) in aqua dest. and 2x DMEM (Merck) in a 1:1 ratio. After 5 days at 37°C the overlay was removed, cells fixed with 1 ml/well of 70 % ethanol for 30 min at room temperature and stained with 1 ml/well of 0.3 % crystal violet (sigma-aldrich) in 20 % methanol for 10 min at room temperature. In addition, for HK/1/68, PR/8/34 and Pan/2007/99 virus stocks used in experiments for 16 hrs infections and confocal microscopy the fluorescent focus-forming units (FFU) had been determined based on nuclear NP signal.

For the rescue of A/Hong Koug/1/1968ΔNS1 (HK/1/68ΔNS1) via reverse genetics the NS1 open reading frame was eliminated from the respective plasmid as described previously (Stech et al. 2008) applying one mega-primer (Sequence: 5’ GAAGTTGGGGGGGAGCAAAAGCAGGGGACAAAGACATAATGGATTCTAACACTGTGTCAAGTTTTCAGGACATACTATTGAGGATGTCAAA AATGCAATTGGGGTCCTCATCGGAGGACTTGAATGGAATGATAACACAGTTCGAGTCTCTAAAACTCTACAGAGATTCGCTTGGGGAAGCAGTAATGAGAATGGGAGACCTCCACTCACTCCAAAACAGAAACGGAAAATGGCGAGAACAGTTAGGTCAAAAGTTCGAAGAGATAAGATGGCTGATTGAAGAAGTGAGACACAGATTGAAGACAACAGAGAATAGTTTTGAGCAAATAACATTTATGCAAGCCTTACAGCTACTATTTGAAGTGGAACAGGAGATAAGAACTTTCTCGTTTCAGCTTATTTAATGATAAAAAACACCCTTGTTTCTACTAATAACCCGGCGG 3’) and the plasmid was checked by sequencing. HK/1/68ΔNS1 was recovered using the altered eight plasmid system on MDCKII NS1 cells.

### Stable cell line generation

A549-HA-TRIM25 and A549-eGFP-IRF3 were generated using lentiviral transduction with pWPI-HA-TRIM25 and pWPI-eGFP-IRF3, respectively, which were generated by Gateway-shuttling from TRIM25 pENTR221 (Invitrogen) into a Gateway-compatible derivative of the pWPI-puro lentiviral vector (Trotard et al. 2015). Briefly, lentiviral particles were produced on HEK293T cells by calcium phosphate transfection with three plasmids at a 3:1:3 ratio: 1) pCMV-ΔR8.91, expressing HIV gag-pol; 2) pMD2.G, encoding the VSV-G glycoprotein; and 3) the respective pWPI construct. pCMV-ΔR8.91, pMD.2G, and pWPI were kind gifts from Prof. Didier Trono (Lausanne, Switzerland). Supernatants were harvested 48, 56 and 72 hours post transfection and filtered with a 40-micron filter for a cell free supernatant containing lentiviral particles and stored at −80°C. A549 cells were seeded in a 24 well plate at 1×10^5^/well and cultured overnight in supernatant containing lentiviral particles. Successfully transduced cells were selected adding 1 μg/ml puromycin to the medium.

### Immunofluorescence

Cells subjected for immunofluorescence were fixed using 4% paraformaldehyde in PBS pH 7.4 for 10 min at room temperature. For permeablisation cells had been treated with 0.5 % Triton X-100 for 5 min. Blocking was performed with 5 % BSA in PBS for 10 – 30 min at room temperature. Immunostaining was performed in a humidified chamber for 1 h at room temperature. Cells had been washed three times with PBS between first and secondary antibody incubation. For co-localization studies employing confocal microscopy the following antibody combinations were used: mouse anti-non-structural protein 1 (Santa Cruz)/goat anti-mouse Alexa568 (ThermoFisher Scientific), rabbit anti-TRIM25 (abcam)/goat anti-rabbit Alexa647 (ThermoFisher Scientific). For determination of infection rates employing wide field fluorescence microscopy the following antibodies in combination with a DAPI counter stain were used: mouse anti-nucleoprotein/goat anti-mouse Alexa488 (ThermoFisher Scientific). Cells grown on cover slips were mounted on microscopy slides using Mowiol.

Combined fluorescence *in situ* hybridization (F*i*sH) and immunofluorescence (IF). F*i*sH on viral RNA (full length-ssRNA molecules) was carried-out as described previously using same F*i*sH probes targeting all vRNA segments simultaneously (Haralampiev et al. 2017). In brief, A549 cells seeded in 3.5 cm MaTek dishes were fixed in 10 % formalin 16 hrs post infection for 10 min at room temperature. A mild permeabilisation was achieved by incubating the cells in 70 % ethanol at 4°C overnight. After rehydration through two washing steps with 2x SSC buffer (2x SSC in RNase-free water, 2 mM VRC) cells were blocked in RNase-free BSA (1:10 BSA in 2x SSC buffer) for 30 min at room temperature. Immunostaining of NS1 and TRIM25 was realized by using the following antibody combination: mouse anti-NS1/goat anti-mouse Alexa 488 (ThermoFisher Scientific) and rabbit anti-TRIM25 (abcam)/goat anti-rabbit Atto647N (sigma). Cells were incubated with first and secondary antibody using a humidified chamber for 1 h at room temperature and washed with 2x SSC buffer three times after each incubation step. F*i*sH on viral RNA was performed by incubating the cells with the hybridization buffer (10 % formamide, 2x SSC, 10 % Dextran-sulfate, 2 mM VRC and 100 nM of each F*i*sH probe set per genome segment in RNase-free water) for 4 hrs at 37°C. Stained samples were washed twice with formamide buffer (10 % formamide, 2x SSC in RNase-free water) for 15 min at room temperature. Microscopy was performed in 2x SSC buffer immediately.

### Widefield microscopy

Image Acquisition was performed using an inverted fluorescence widefield microscope (Olympus IX70) and a 20x/0.40 air LCPlanFL objective. Single planes were measured in the DAPI channel and the GFP (Alexa488) channel.

### Confocal microscopy

Image acquisition was performed using an inverted fluorescence confocal microscope (Leica SP8) and a 63x/1.40 oil immersion objective. Images are shown as single planes with a pixel size of 120 nm. Maximal projections of z-stacks were used for data analysis of NS1 and TRIM25 localization.

### STED nanoscopy

Super-resolution microscopy on FisH/IF samples was performed using a Two-color-STED microscope(Abberior Instruments GmbH). For confocal and STED microscopy a 100x Olympus UPlanSApo (NA 1.4) oil immersion objective was used. For excitation (λ = 594 nm and λ = 641) nm) a nominal laser power of 20 % and for STED (λ = 775 nm, max. power = 1.2 W) a nominal laser power of 70 % was applied. Pixel size was set to 60 nm (confocal) and 15 nm (non-diffracted), respectively. Minor adjustments of contrast and brightness of acquired images as well as Richardson-Lucy deconvolution with a regularisation parameter of 0.001 (stopped after 30 iterations) were carried out using Imspector software (Abberior Instruments GmbH).

### Data analysis

Infection rates were calculated by determining the proportion of cells being positive for nuclear nucleoprotein (NP) localization at indicated times (hours post infection) based on widefield microscopy images. Total cell number was defined by DAPI counterstaining of nuclei and thresholded mean NP signal intensity in the nuclei (= DAPI positive ROIs) were quantified automatically using image J software. In brief, a cell was considered being infected fulfilling the following parameters: (i) more than 80 % of the nuclear area is covered by NP signal and (ii) NP signal intensity was above 20/255 for A549 wt and A549-eGFP-IRF3, and 100/255 for A549-HA-TRIM25, repectively. For noise reduction, Gaussian filtering (sigma = 2) was applied to raw images. For quantitative analysis of microscopy data a semi-automated approach using Image J software and R program was applied. Batch analysis was performed for all images with a pixel size of 120 nm. A maximum projection of acquired z-stacks for each channel separately was used as starting data set. For background correction the mean intensity of background area of at least five images per dataset was calculated and subtracted for each image, respectively. Cell boundaries were drawn as ROI manually and corrected by an adjustable watershed function. Further, a mask (A) of these ROIs was created. DAPI signal was used to create a mask (B) of nuclei by thresholding (80/255) and manual correction. Mask A and B were subtracted representing the cytosolic regions (= mask C). For segmentation of NS1 aggregates, mask C was subtracted from the NS1 image and then a Gaussian blur function (sigma = 2) was performed to reduce noise. Threshold was set to 180/255. After running an adjustable watershed function (tolerance = 0.3), the particle size was limited to 0.6 μm^2^ - infinity creating mask D, which enabled counting of aggregates per cell. Analysis of signal distribution of NS1 and TRIM25 at aggregates was performed as follows: An additional mask E representing the cytosol without the area occupied by aggregates was created by inverting the sum of mask C and mask D. For each cell (= ROI) the mean intensity of NS1 and TRIM25 at aggregates (= mask D) and the surrounding cytosol area (= mask E) was measured. Furthermore, the mean intensity of the IRF3-GFP signal at the cytosol as well as the mean intensity of the NS1 signal of entire cells was measured. These values were combined by using the R program resulting in a tabular presentation of all obtained information. Cells had been assigned “infected” when mean NS1 intensity value was above 40. Only cells exhibiting more than three NS1 aggregates were included into the analysis. For RNA localization analysis at TRIM25 positive NS1 aggregates, NS1 aggregates were segmented as described before with a threshold of 70/255. Only aggregates in the cytosolic area were considered for measurements of TRIM25 and RNA signal on NS1 aggregates. Aggregates with an integrated mean fluorescence intensity above 10 were considered positive for either TRIM25 and/or RNA.

### Co-immunoprecipitation

A549-HA-TRIM25 cells were infected with HK/1/68, Pan/2007/99, and PR/8/34 (MOI = 1). 16 hrs post infection cells were lysed for 30 min gently shaking on ice with IP lysis buffer (20 mM TRIS (pH7.6), 100 mM NaCl, 1 % NP40, 50 mM NaF) supplemented with protease inhibitor cocktail (Roche). Lysates were centrifuged for 30 min at 14000 rpm and 4°C. An aliquot of the postnuclear fraction were kept as input control; the remaining lysate was mixed with 20 μl HA beads (Sigma Aldrich) which were prior washed three times with IP lysis buffer. Beads and lysates were incubated rotating for 4 hrs at 4°C before they were washed three times with IP lysis buffer for 15 min at 4°C. Beads were additionally washed two times with ice cold PBS. HA tagged proteins were eluted twice with 100 μl 5 % SDS in PBS shaking for 5 min at 1100 rpm followed by a 14000 rpm centrifugation. Supernatant, containing the eluted proteins were precipitated using acetone. Input and pulldown samples were separated on a 12 % SDS-PAGE and transferred onto PVDF membrane. Specific proteins were detected using anti-TRIM25 (BD biosciences), anti-NS1 (Santa Cruz Biotechnology), and anti-beta-actin (Sigma Aldrich) antibodies.

### Immunoblotting

Whole-cell lysates were prepared using 6x lysis buffer (9.75 ml Tris 0.5 M (pH6.8), 15 ml glycerol, 15 ml 10 % SDS, 3.75 ml beta-mercaptoethanol, 30 mg bromophenol blue, 6.5 ml H2O). Protein samples were resolved by 8 % SDS-PAGE at 120 Volt for 2 hrs. Then proteins were transferred onto PVDF membranes (Biorad). Membranes were dried and re-hydrated in methanol, washed twice for 10 min in PBS at room temperature, and probed with first antibodies (rabbit anti-phospho-IRF3, catalogue no.4947, Cell Signalling, mouse anti-NS1, catalogue no. sc-130568, Santa Cruz, rabbit anti-calnexin, catalogue no. ADI-SPA-865-F, Enzo Life Science, and rabbit anti-IRF3, catalogue no. 11904S, Cell Signalling) diluted 1:1000 in 5 % BSA in 1xTBST at 4°C overnight. After antibody incubation, membranes were washed three times in 1xTBST (anti-phospho-IRF3) or 1xPBST (anti-IRF3, anti-NS1, anti-calnexin) for 10 min. Membranes were incubated with secondary goat anti-rabbit HRP antibody (sigma-aldrich) or secondary goat anti-mouse HRP antibody (sigma-aldrich) for 1 h at room temperature. Protein bands were visualized using the ECL ChemCam imager 3.2 (Intas). Relative protein expression was determined using 1D analysis software (Intas).

## 3. Results

### 3.1. Host TRIM25 accumulates in viral NS1 aggregates in infected cells in late stages of infection, in a strain-dependent manner

In our study, we focused on the interaction of cellular antiviral factors and the non-structural IAV protein 1 (NS1), which opposes their function. To assess whether cells were activated via the antiviral interferon pathway, we introduced stable expression of eGFP-IRF3 in A549 cells using lentiviral vector transduction and subsequent puromycin selection. The introduction of eGFP-IRF3 into A594 cells may affect the biophysical status of the cell and thus, cellular processes and IAV infection. To validate the infection kinetics, we examined the protein level of phosphorylated IRF3 as a marker for interferon signaling and activation of antiviral defense. Additionally, we monitored NS1 as a marker for infection progression. Immunoblotting of whole cell lysates of A/Puerto Rico/8/1934 (PR/8/34) infected A549 wt cells and A549-eGFP-IRF3 cells at different times post infection (pi) showed a reduced rate of infection progression in A549-eGFP-IRF3 cells compared to A549 wt cells, but a fairly equal absolute abundance of NS1 at 16 hrs pi (Figure 1A and B, Figure S1 A for a comparison of infection rates in A549 wt and A549-eGFP-IRF3). Due to overexpression of eGFP-IRF3, elevated levels of phosphorylated IRF3 are detected in A549-eGFP-IRF3 cells (Figure 1 A, C).

**Figure 1.**
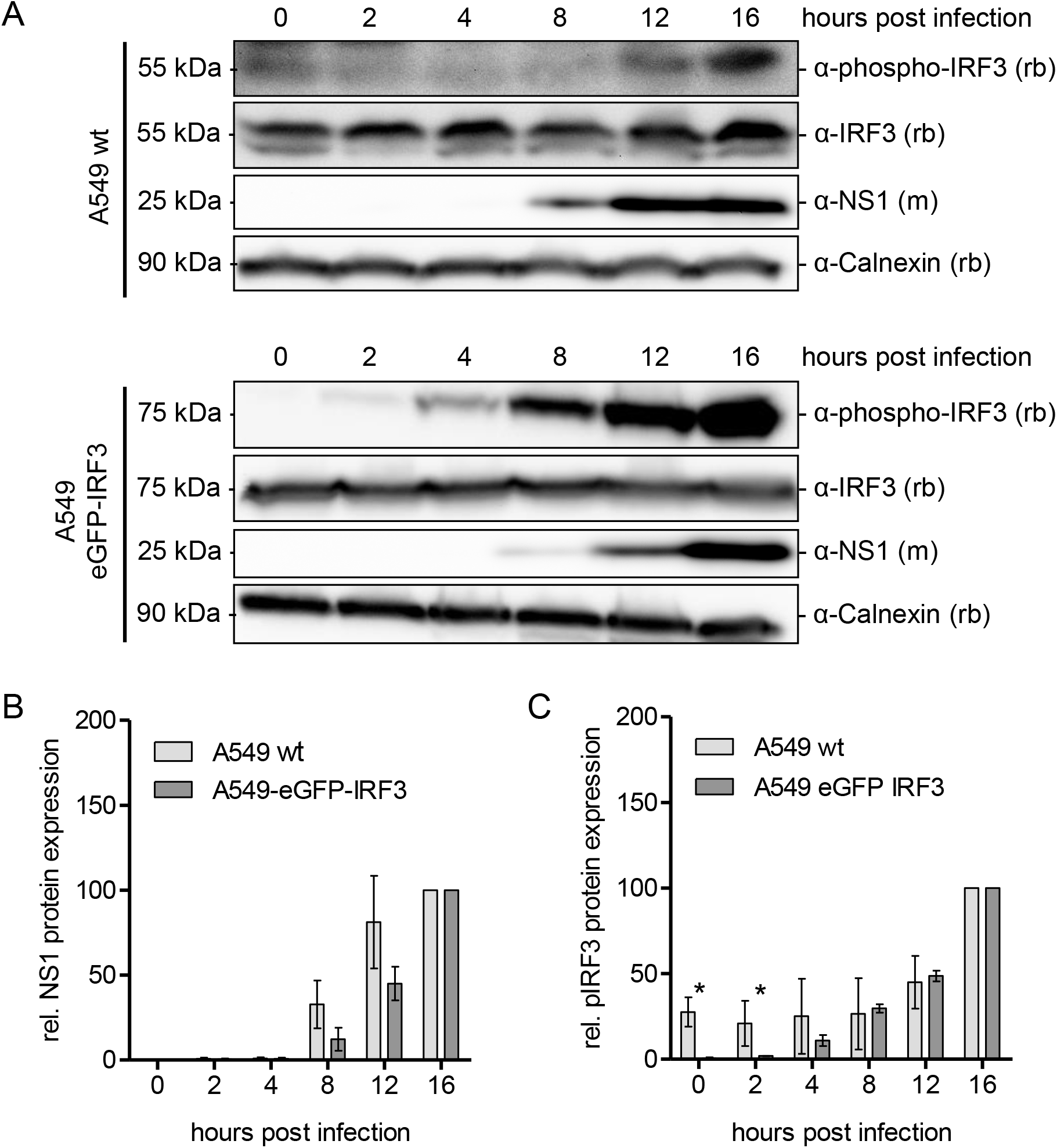
Influenza A virus non-structural protein 1 level are comparable in A549 wt and A549-eGFP-IRF3 cells. **(A)** Immunoblot analysis of phosphorylated IRF3 (phospho-IRF3) and non-structural protein 1 (NS1) levels in whole-cell lysates from non-infected (0) and PR/8/34 infected A549 wildtype and A549-eGFP-IRF3 cells (times post infection as indicated). Calnexin was used as loading control. Panels are derived from a single cropped blot with lanes 1–6 belonging to the same membrane and from the same experiment for A549 wt and A549-eGFP-IRF3, respectively. Of note, A549 wt and A549-eGFP-IRF3 cells vary in IRF3 and rGFP-IRF3 levels, respectively, therefore the blots cannot be used for quantitative comparison between the cell lines. To detect phosho-IRF3 longer exposure times were applied. **(B)** Relative NS1 protein expression in A549 wildtype and A549-eGFP-IRF3 cells normalized to calnexin of three independent experiments. Mean values are plotted as mean (SD). The results for the cell lines are not significantly different from each other. **(C)** Relative phosphorylated IRF3 protein expression in A549 wildtype and A549-eGFP-IRF3 cells normalized to calnexin of two independent experiments. Values are plotted as mean (SD). (* p < 0.05) For each cell type normalized protein levels at 16 hours post infection were set to 100 %.

To investigate the strain dependence of NS1 interaction with TRIM25, we infected A549-eGFP-IRF3 cells with three different IAV strains: A/Puerto Rico/8/1934 (PR/8/34, H1N1), A/Hong Kong/1/1968 (HK/1/68, H3N2) and A/Panama/2007/1999 (Pan/2007/99, H3N2). After 16 hrs pi, cells were immunostained with anti-NP antibodies and the nucleus was counterstained with DAPI. The infection rate for IAV- and mock-infected A549-eGFP-IRF3 cells was calculated by determining the number of cells with nuclear NP in relation to the total number of cells present based on fluorescent widefield microscopy images (Figure S1 B).

The results for all used IAV strains showed values between about 30 and 50 % of infected cells. In a second step, simultaneously infected A549-eGFP-IRF3 cells were immunostained for IAV NS1 and TRIM25. 500 nm-z-stacks were recorded with an inverse confocal laser scanning microscope and quantitative data analysis was performed on maximum projections. Representative single layer images show cytosolic NS1 aggregates at 16 hrs pi in PR/8/34, HK/1/68, and Pan/2007/99 infected cells (Figure 2 A, white arrowheads). By initial visual observation we found a low frequency of these aggregates in PR/8/34 infected cells compared to HK/1/68 and Pan/2007/99 infected cells. For the quantitative analysis of the intracellular localization of NS1 and TRIM25 we used a semi-automated image analysis approach with self-written algorithms (software R). We generated masks specific for the whole cell volume, only the nucleus and NS1 aggregates to distinguish between these regions (Figure S2). First, we quantified the occurrence of NS1 aggregates in infected cells (Figure 2 B). The proportion of cells with NS1 aggregates is significantly increased in Pan/2007/99 and HK/1/68 compared to PR/8/34 infected cells (Figure 2 B). It is noteworthy that less than 5 % of cells infected with PR/8/34 have NS1 aggregates. In HK/1/68 and Pan/2007/99 infected cells there are 4 to 6 times higher NS1 aggregation levels compared to H1N1 infected cells. Despite the low frequency of NS1 aggregates in PR/8/34 infected cells, the analysis of the TRIM25 distribution on aggregates compared to extranuclear space showed that on NS1 aggregates about twice the amount of TRIM25 is found (Figure 2 C). According to the frequency of NS1 aggregates in HK/1/68 and Pan/2007/99 infected cells, there are significantly higher amounts of TRIM25 that colocalize to NS1 aggregates. In summary, TRIM25 accumulates at 16 hrs pi on NS1 aggregates for all three IAV strains applied, although there are disproportionately large amounts of TRIM25 on NS1 aggregates in PR/8/34 infected cells.

**Figure 2.**
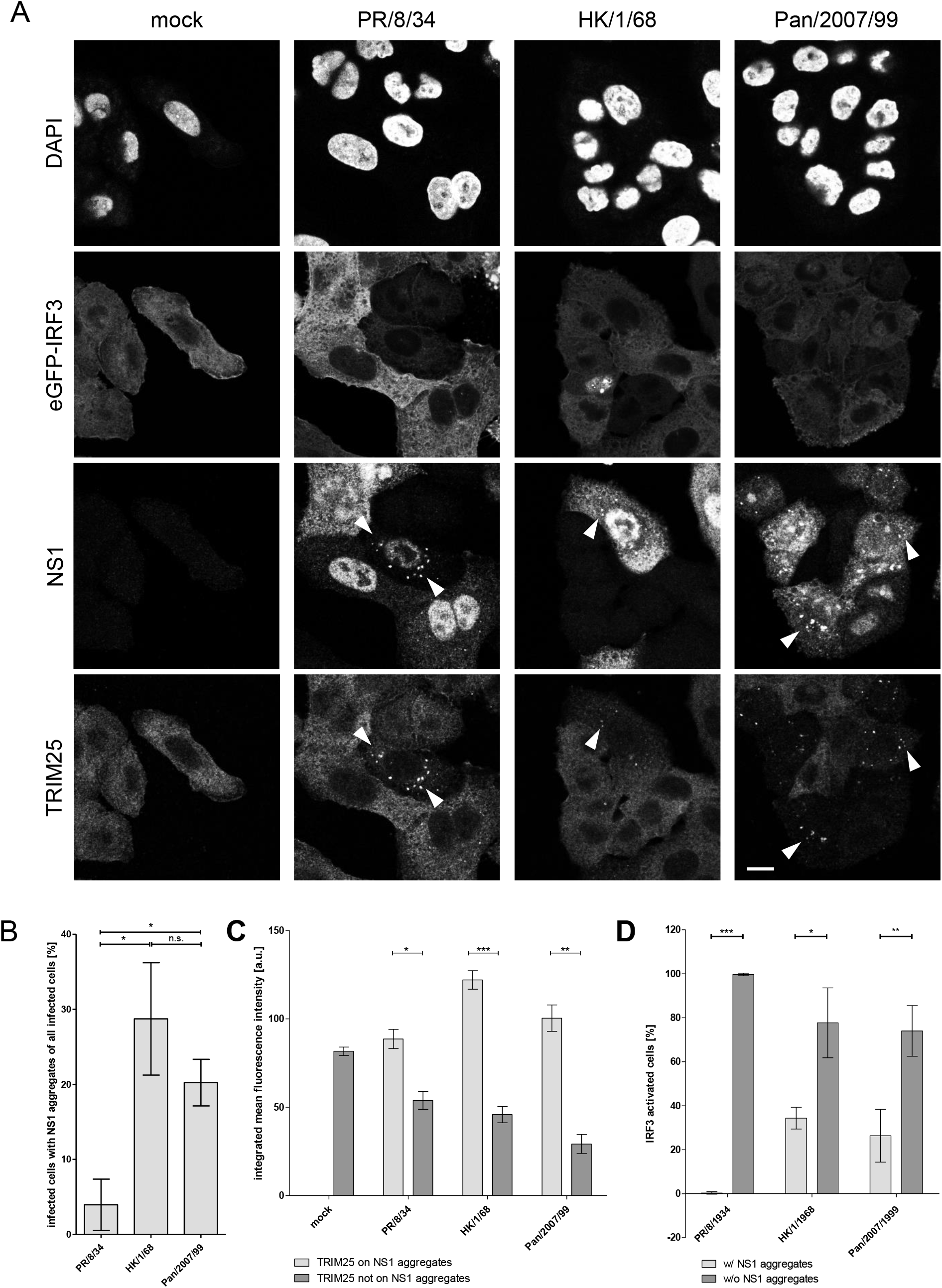
Abundance of NS1 cytosolic aggregates is significantly enhanced in A/Pan/2007/99 infected A549-eGFP-IRF3 cells 16 hours post infection and TRIM25 is localized to cytosolic NS1 aggregates resulting in diminished IRF3 activation. **(A)** A549-eGFP-IRF3 cells subjected for immunofluorescence were mock, PR/8/34, HK/1/68, and Pan/2007/99 infected for 16 hours (MOI = 1). Immunostaining was performed using the following first and secondary antibody combinations: mouse anti-NS1/goat anti-mouse Alexa568 and rabbit anti-TRIM25/goat anti-rabbit Alexa647. White arrowheads indicate sites of NS1 aggregates with colocalizing TRIM25 signals in IAV infected cells. Image acquisition was performed using an inverted confocal microscope. Images are shown as single planes with a pixel size of 120 nm. Scale bar corresponds to 20 μm. Quantitative analysis is based on maximal projections of 500 nm z-stacks. **(B)** Graph shows the frequency of cytosolic NS1 aggregates in PR/8/34, HK/1/68, and Pan/2007/99 infected A549-eGFP-IRF3 cells. The abundance of NS1 aggregates was analyzed based on NS1 signals in the extra-nuclear space. Results represent the mean (SD) of three independent experiments (n: HK/1/68 = 292 cells, Pan/2007/99 = 217 cells, PR/8/34 = 265 cells). **(C)** Integrated (= per selected area, please refer to “data analysis” section for further information) mean fluorescence intensity of TRIM25 was analyzed with respect to cytosolic NS1 aggregates. Results represent the mean (SD) of three independent experiments (n: HK/1/68 = 80 cells, Pan/2007/99 = 44 cells, PR/8/34 = 11 cells, mock = 57 cells). Propagation of uncertainty was included in statistical analyses. **(D)** Graph shows IRF3 nuclear translocation frequency in cells with and without NS1 aggregates. Quantitative analysis was performed for NS1 aggregates with respect to nuclear IRF3 signal. Results represent the mean (SD) of three independent experiments.

### 3.2. A549-eGFP-IRF3 cells with cytosolic NS1 aggregates show decrease nuclear IRF3 signal

Next, we attempt to answer whether NS1 cytoplasmic aggregates interfere with the activation of antiviral defense. To this end, we analyzed the correlation between NS1 aggregate frequency and nuclear localization of activated IRF3. By monitoring the IRF3 translocation into the nucleus, we were able to identify cells in which antiviral signaling was triggered (Figure S3). Interestingly, our analysis revealed in PR/8/34 infected cells that the majority of cells with cytosolic NS1 aggregates having no IRF3 activation (Figure 2 D), suggesting an effective block of interferon I induction. While in Pan/2007/99 and HK/1/68 infected cells this phenotype is less pronounced, in both cases about 30 % of the IRF3 activated cells have cytosolic NS1 aggregates. For the H3N2 variants studied, we observed a milder correlation between the occurrence of cytosolic NS1 aggregates and the block of IRF3 activation. However, we found significantly less activation of antiviral signaling by a significantly reduced frequency of IRF3 activation among cells harboring cytosolic NS1 aggregate.

### 3.3 NS1 is co-precipitated with TRIM25 from IAV-infected A549-HA-TRIM25 cells

To validate the results of our imaging approach, we have analyzed potential NS1 TRIM25 interactions using coimmunoprecipitation. To this end, we infected A549-HA-TRIM25 cells with PR/8/34, HK/1/68, and Pan/2007/99 and their respective ΔNS1 variants to verify assay specificity in NS1 detection. Infection was determined by western blotting of whole cell lysates (Figure S4 A) and determination of infection kinetics based on nuclear NP signal quantification (Figure S4 B). We found reduced levels of M1 produced in cells infected with ΔNS1 variants in comparison to the respective wt strain.

Furthermore, the cells were lysed after 16 hrs pi and HA-TRIM25 immunoprecipitated with anti-HA antibodies. Western blotting for NS1 showed robust co-precipitation of NS1 with HA-TRIM25 for all investigated IAV strains (Figure 3). In mock, PR/8/34 ΔNS1, HK/1/68 ΔNS1, and Pan/2007/99 ΔNS1 infected A549-HA-TRIM25 cells, and infected A549 wt cells no signal for NS1 was detected. The pull-down NS1 values in HK/1/68 and Pan/2007/99 infected cells appear to be elevated compared to the situation in PR/8/34 infected cells (Figure 3 lower panel). Please remember that in PR/8/34 infected cells the amount of NS1 aggregates was reduced (Figure 2 B).

**Figure 3.**
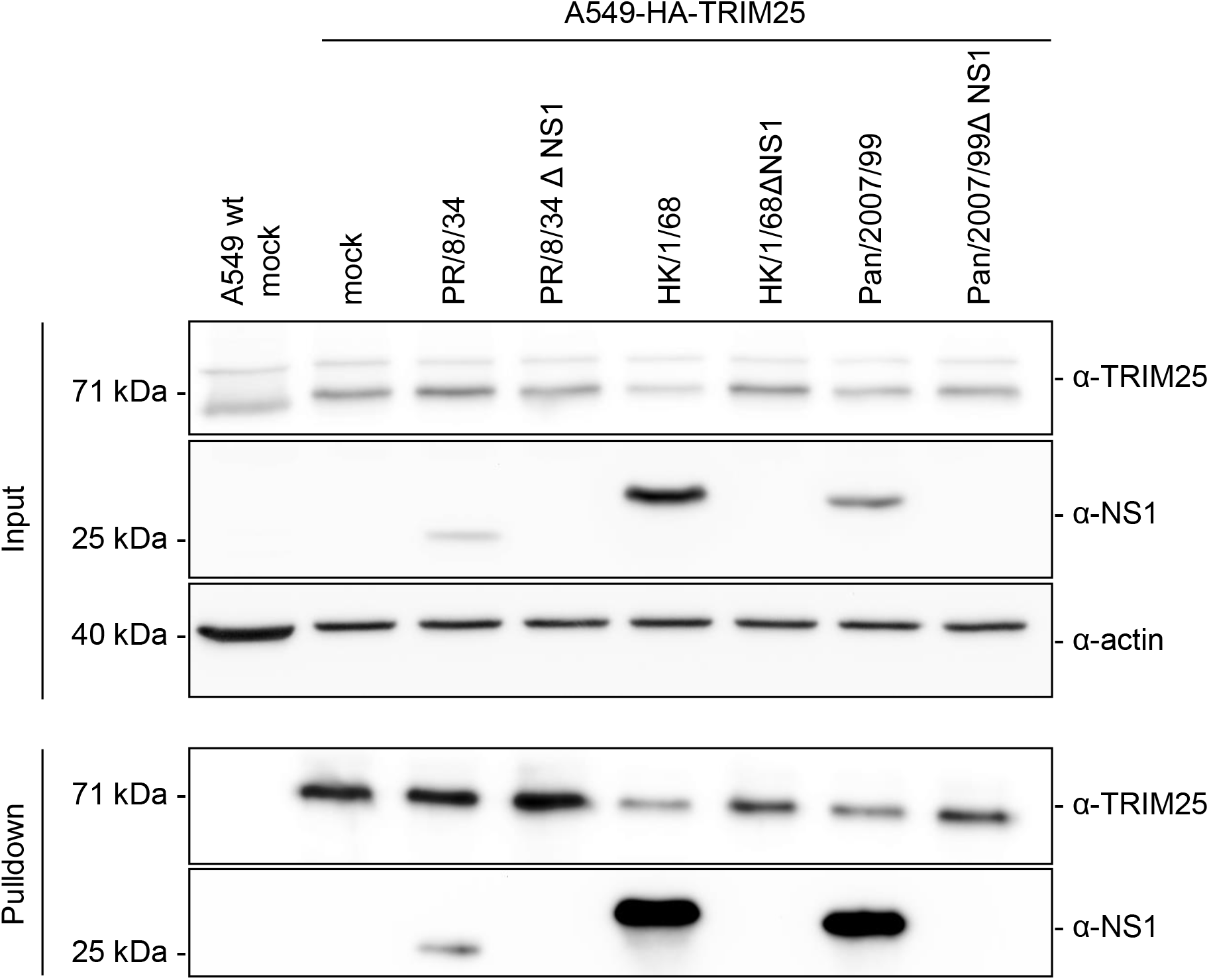
IAV non-structural protein 1 is co-immunoprecipitated with HA-TRIM25 using anti-HA antibody in A549-HA-TRIM25 cells. A549-HA-TRIM25 cells were infected with PR/8/34, HK/1/68, and Pan/2007/99 and their respective ΔNS1 variants. Whole cell lysates were prepared 16 hours post infection. Mock infected control and ΔNS1 IAV variants show no unspecific NS1 signals. Actin was used as loading control. Panels for the input and the pulldown are derived from a single cropped blot with lanes 1-8 belonging to the same membrane and from the same experiment, respectively.

We observed comparable TRIM25 values in A549 wt cells and A549-HA-TRIM25 cells. In addition, the TRIM25 level does not change during IAV infection in A549-HA-TRIM25 cells (Figure 3). We have to admit that we cannot control the accessibility of antibody binding sites, but the preparation protocol includes a denaturing SDS treatment. Interestingly, we found the mean intensity of NS1 signals in cells with aggregates is increased compared to cells without NS1 aggregates for all three tested IAV wt strains which might argue for a dose dependent NS1 cytosolic aggregation (Figure S5 A). Generally, the mean intensity of TRIM25 signal in the nuclear and extranuclear region is not significantly different in PR/8/34 infected A549-eGFP-IRF3 cells compared to those infected with HK/1/68 or Pan/2007/99 (Figure S5 B).

### 3.4. A minor portion of TRIM25-NS1 complexes is positive for viral RNA

Assuming that the binding of viral RNA (vRNA) is important for the formation of antiviral complexes, we followed this hypothesis by investigating the presence of IAV vRNA in TRIM25-NS1 cytosolic complexes using superresolution microscopy (STED nanoscopy). For the simultaneous visualization of IAV vRNA and TRIM25 in nanoscale resolution, we have developed an approach that combines RNA *in situ* hybridization (RNA F*i*sH) and protein detection by immunofluorescence. Atto594 labeled IAV vRNA F*i*sH probes specific for Pan/2007/99 were kindly provided by Andreas Herrmann, Humboldt University, Berlin, Germany (Haralampiev et al. 2017).To investigate a possible colocalization of TRIM25 and vRNA with NS1, a diffraction-limited channel in confocal imaging mode was added to the analysis (Figure 4). Nonspecific signal induced by vRNA F*i*sH or immunolabeling of NS1 were negligible or absent, as shown in mock and Pan/2007/99ΔNS1 infected A549 wt cells. Therefore, we assume a high specificity of the combined F*i*sH/IF staining. In Pan/2007/99 ΔNS1 infected A549 wt cells, no accumulation of TRIM25 was detected in the cytosol, suggesting NS1-triggered localization of TRIM25 at sites with elevated NS1 levels. In accordance with the results of Meyerson and colleagues (Meyerson et al. 2017) strong signals of TRIM25 colocalizing on vRNA were found in the nucleus of Pan/2007/99 ΔNS1 infected A549 cells. Here, TRIM25 cannot be sequestered by NS1 in the cytosol and thus, translocates into the nucleus to impede viral RNA elongation as previously described [30]. By contrast, we again observed TRIM25 accumulation on NS1 cytosolic aggregates in Pan/2007/99 wt infected A549 cells (Figure 4, white arrowheads). Only 15 % out of 143 analysed TRIM25-NS1 aggregates were positive for vRNA, suggesting that vRNA is not a mandatory factor for the interaction of NS1 with TRIM25 to form cytosolic aggregates at late stage of infection.

**Figure 4.**
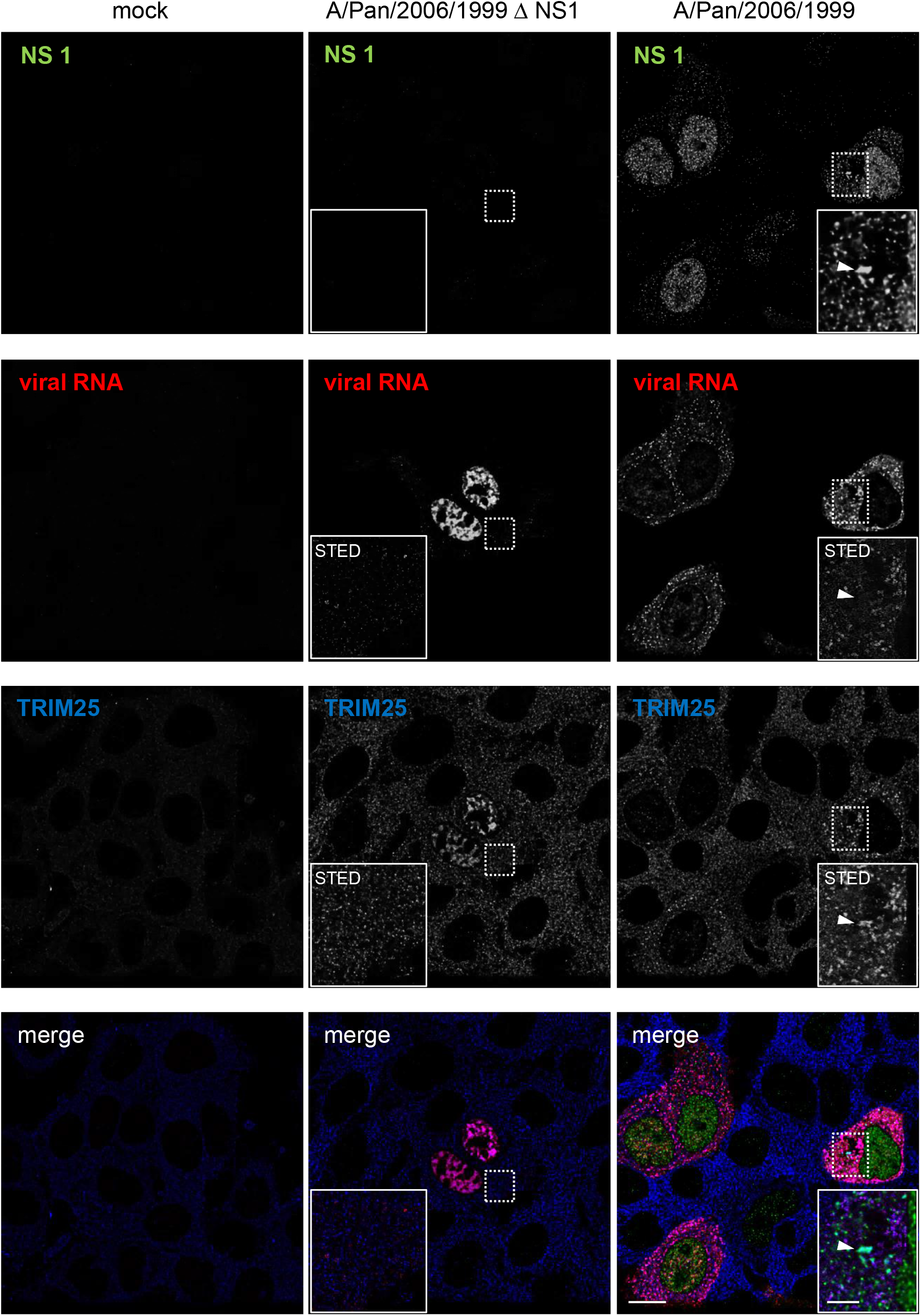
TRIM25 is located in NS1 aggregates but viral RNA does not accumulate at these sites. A549 wt cells were mock, Pan/2007/99, and Pan/2007/99 ΔNS1 infected for 16 hours (MOI = 1). Arrowheads point at sites of NS1 aggregates with TRIM25 colocalization. Images are shown as single planes and image acquisition was performed using 60 nm (= confocal mode, NS1) and 15 nm (= STED, viral RNA and TRIM25 signal) pixel size. Dashed square indicates zoom-in area. Scale bars correspond to 10 μm for overview and 3 μm for zoom-in images, respectively.

## 4. Discussion

The study described herein elucidates the scavenger effect of NS1 on cytosolic TRIM25 to interfere with its antiviral function in the nucleus. We found a strain-specific manifestation of NS1 aggregation in the cytosol. Previous studies suggest that the binding of NS1 to TRIM25 is a prerequisite for blocking IRF3 activation and TRIM25 localization at the nucleus (Kuo et al. 2010). Accordingly, we found lower concentrations of activated IRF3 in cells with cytosolic NS1 aggregates. The basal level of phosphorylated IRF3 is increased in A549-eGFP-IRF3 cells compared to wild-type cells, while NS1 expression appears at 16 hrs pi in equal amounts as in wild-type cells. Thus, the influence of NS1 on IRF3 in our system may be underestimated. Single cell analysis allowed us to distinguish exactly between the phenotypes identified: cells with equally distributed NS1 in the cytosol, cells with strong cytosolic NS1 aggregates, and cells without NS1 signal. For decades, strain dependency has been of great relevance in IAV biology research. To address this issue, we have included a well characterized H1N1 and two H3N2 pandemic strains in our study to clarify whether they perform similar mechanisms to block antiviral signaling pathways. In fact, we confirm strain variations in the ability to inhibit the activation of IRF3. Using fluorescence imaging, we found the highest proportion of cells with NS1 aggregates in HK/1/68 infected cells and the strongest TRIM25 localization at these sites at 16 hrs pi. It is remarkable that in PR/8/34 infected cells the proportion of cells with cytosolic NS1 aggregates is 4 to 6 times lower than in HK/1/68 or Pan/2007/99 infected cells, while in PR/8/34 infected cells similar amounts of TRIM25 are localized in cytosolic NS1 aggregates. This finding is supported by co-immunoprecipitation of TRIM25-HA and NS1: in the case of PR/8/34 infected cells, higher levels of TRIM25-HA appear in the pulldown, while lower levels of NS1 are observed compared to HK/1/68 and Pan/2007/99 infected cells. For both H3N2 strains, higher NS1 values were found in the pulldown, followed by lower TRIM25 levels. For PR/8/34 infected cells, our data showed a strong correlation between the lack of cytosolic NS1 aggregates and the occurrence of nuclear IRF3. Or *vice versa*, we found almost all cells with cytosolic NS1 aggregates without IRF3 nuclear translocation. While H3N2 IAV strains show approx. 30-40 % of IRF3 activated cells with cytosolic NS1 aggregates. Meyerson et al. describe the co-localization of viral RNA and TRIM25 in the nucleus of infected cells (Kuo et al. 2010). We have also observed this phenomenon in Pan/2007/99 ΔNS1 infected cells. To determine whether viral RNA is part of the TRIM25-NS1 cytosolic complex, we performed super-resolution STED microscopy for localization accuracy matters. We used fluorescent probes targeting Pan/2007/99 viral RNA molecules (kindly provided by Andreas Herrmann, University of Berlin, Germany). We also combined viral RNA-F*i*sH with immunofluorescence labelling of proteins for nanoscale imaging. We observed co-localization of TRIM25 and NS1 in Pan/2007/99 infected cells as described above, but only in a minor portion of TRIM25/NS1 double positive aggregates we were able to detect viral RNA in Pan/2007/99 infected A549 cells. However, our results are limited to observations in fixed samples and do not refer to dynamic processes or kinetics. Hence, studies to follow the antiviral complex formation and viral RNA movement in living infected cells in time and space are mandatory.

## 5. Conclusion

Our study addresses the relevance of IAV strain specific variation in blocking cellular antiviral strategies. We found varying levels of TRIM25 on NS1 cytosolic aggregates which might correlate with the capacity to arrest TRIM25 in the cytosol. In addition, vRNA molecules seem to be not important for cytosolic TRIM25/NS1 complex formation.

## Acknowledgments

We are grateful to Andreas Herrmann (Humboldt-Universität zu Berlin, Berlin, Germany), Hans-Georg Kräusslich (University Hospital, Heidelberg, Germany), Marco Binder (German Cancer Research Center, Heidelberg, Germany), and Georg Kochs (University Hospital, Freiburg, Germany) for helpful discussions. We thank, Catharina Gandor and Dominique Dörr (student assistance) as well as Jamie Frankish, Joschka Willemsen, and Sandra Bastian (German Cancer Research Center, Heidelberg, Germany) and Vibor Laketa (IDIP, Heidelberg, Germany) for excellent technical assistance, Sara Dusel (medical student) for support in imaging data acquisition, Jürgen Stech (Friedrich-Loeffler-Institute, Insel Riems, Germany) for the recovery plasmid system of influenza A/Hong Kong/1/1968 and A/Puerto Rico/8/1934, Thorsten Wolff (Robert Koch Institute, Berlin, Germany) for A/Panama/2007/1999 and A/Panama/2007/1999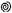NS1, Adolfo Garcia-Sastre (CEIRS program (NIAD Centers of Excellence for Influenza Research and Surveillance), Icahn School of Medicine at Mount Sinai, U.S.A.)) for A/Puerto Rico/8/1934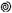NS1, Richard Edward Randall (School of Biology, University of St. Andrews, United Kingdom of Great Britain) for MDCKII-NS1 cell line, and Marco Binder (German Cancer Research Center, Heidelberg, Germany) for A549-eGFP-IRF3 cell line.

## Author Contributions

Anne Weiß: Investigation, Formal analysis, Writing – original draft preparation. Ivan Haralampiev: Design and synthesis of viral RNA F*i*sH probes, Writing – Reviewing and Editing. Susann Kummer: Resources, Conceptualization, Investigation – F*i*sH/IF staining and STED nanoscopy, Project administration, Writing – Reviewing and Editing.

---

**Figure S1.**
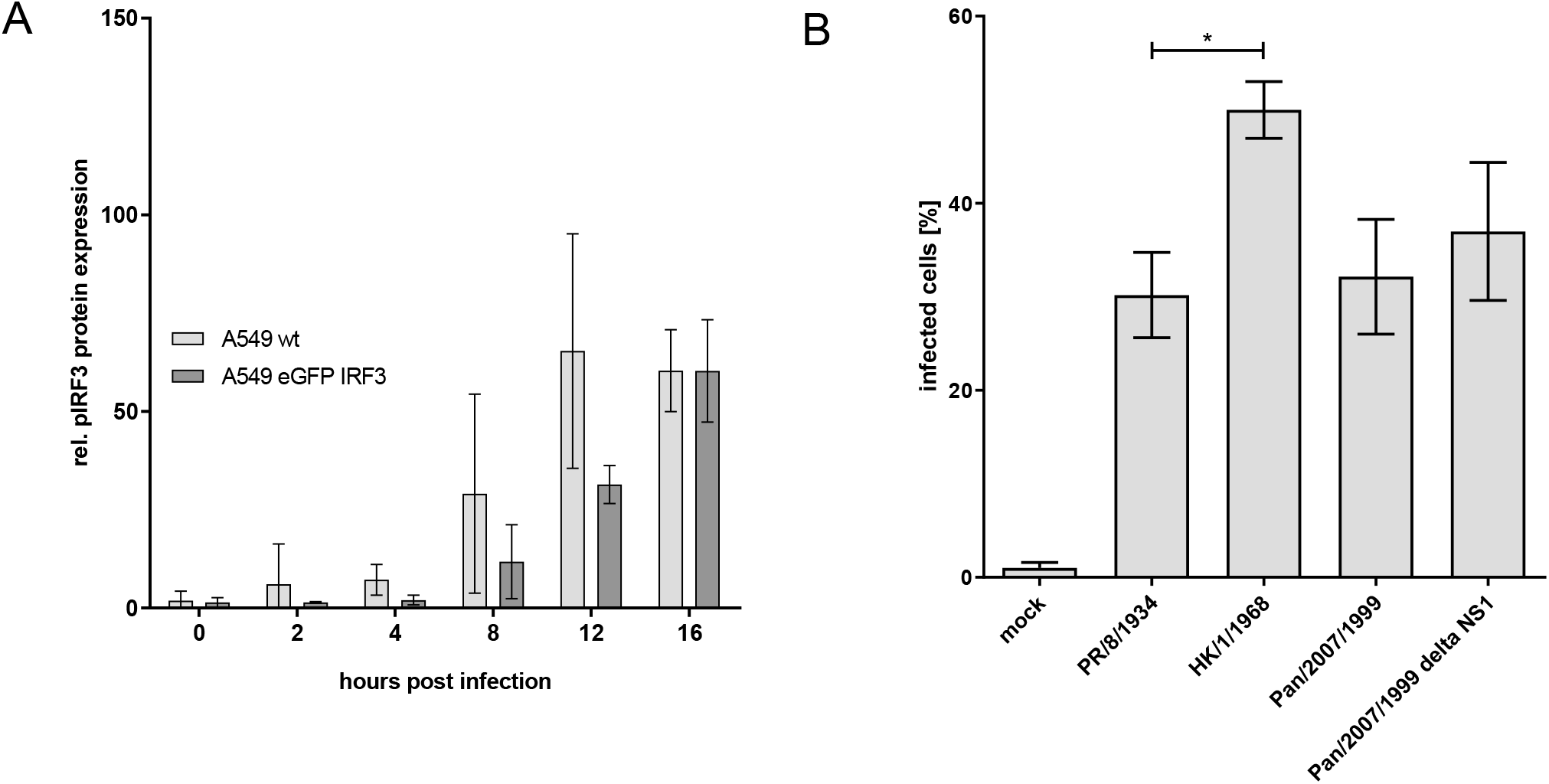
Influenza A virus infection rates are comparable in A549 wt and A549-eGFP-IRF3 cells and infection rates differ between selected IAV strains in A549-eGFP-IRF3 cells. **(A)** A549 wt and A549-eGFP-IRF3 cells subjected for immunofluorescence were mock and PR/8/34 infected for indicated times (MOI = 1). Fluorescence images were acquired using a wide-field microscope. Infection rate was calculated for each cell line by determining the total cell number based on nucleus counterstaining and the proportion of infected cells being positive for nuclear NP applying a semiautomated analysis approach. Graph represents the mean (SD) of three independent experiments (n > 300). Infection rates for A549 wt and A549-eGFP-IRF3 cells are non-significant different from each other, (n: A549 wt, 0 hrs = 1281 cells, 2 hrs = 1142 cells, 4 hrs = 1057 cells, 8 hrs = 845 cells, 12 hrs = 1225 cells, 16 hrs = 1592 cells; A549eGFP-IRF3, 0 hrs = 365 cells, 2 hrs = 713 cells, 4 hrs = 620 cells, 8 hrs = 615 cells, 12 hrs = 539 cells, 16 hrs = 801 cells). **(B)** A549-eGFP-IRF3 cells subjected for immunofluorescence were mock, PR/8/34, HK/1/68 and Pan/2007/99 infected for 16 hours (MOI = 1). Fluorescence images were acquired using a wide-field microscope. Infection rate was calculated for each IAV strain by determining the total cell number based on nucleus counterstaining and the proportion of infected cells being positive for nuclear NP applying a semi-automated analysis approach. Graph represents the mean (SD) of three independent experiments (n > 300). (p < 0.05) Infection rates for Pan/2007/99 is non-significant different from A/Puerto Rico/8/1934.

**Figure S2.**
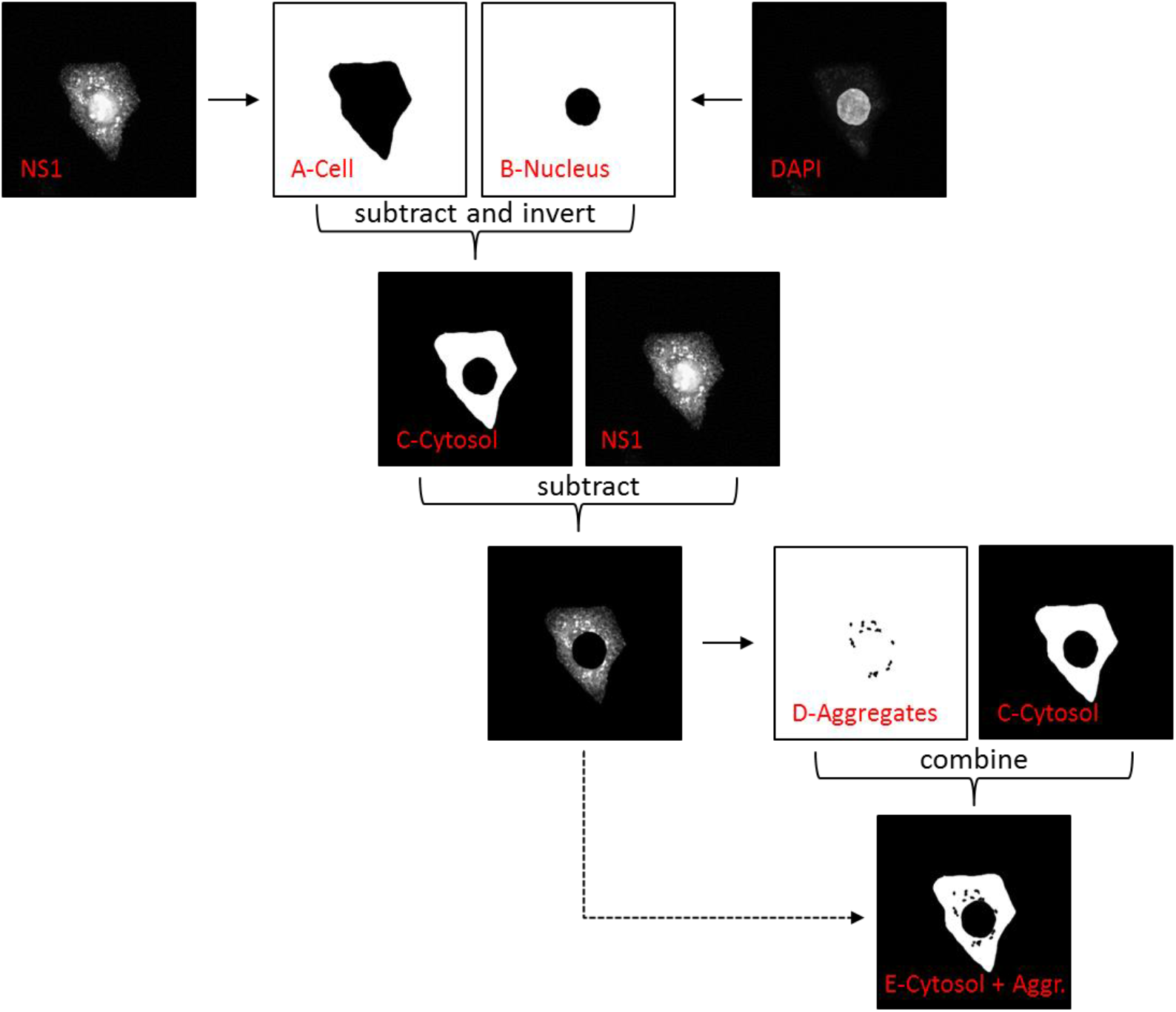
Image analysis was performed by a semi-automated approach using self-written algorithms in software R. The scheme depicts the main steps of the image analysis where the calculations are presented as arrows. Masks are named with letters and respective region. (For explanation please refer to methods section.)

**Figure S3.**
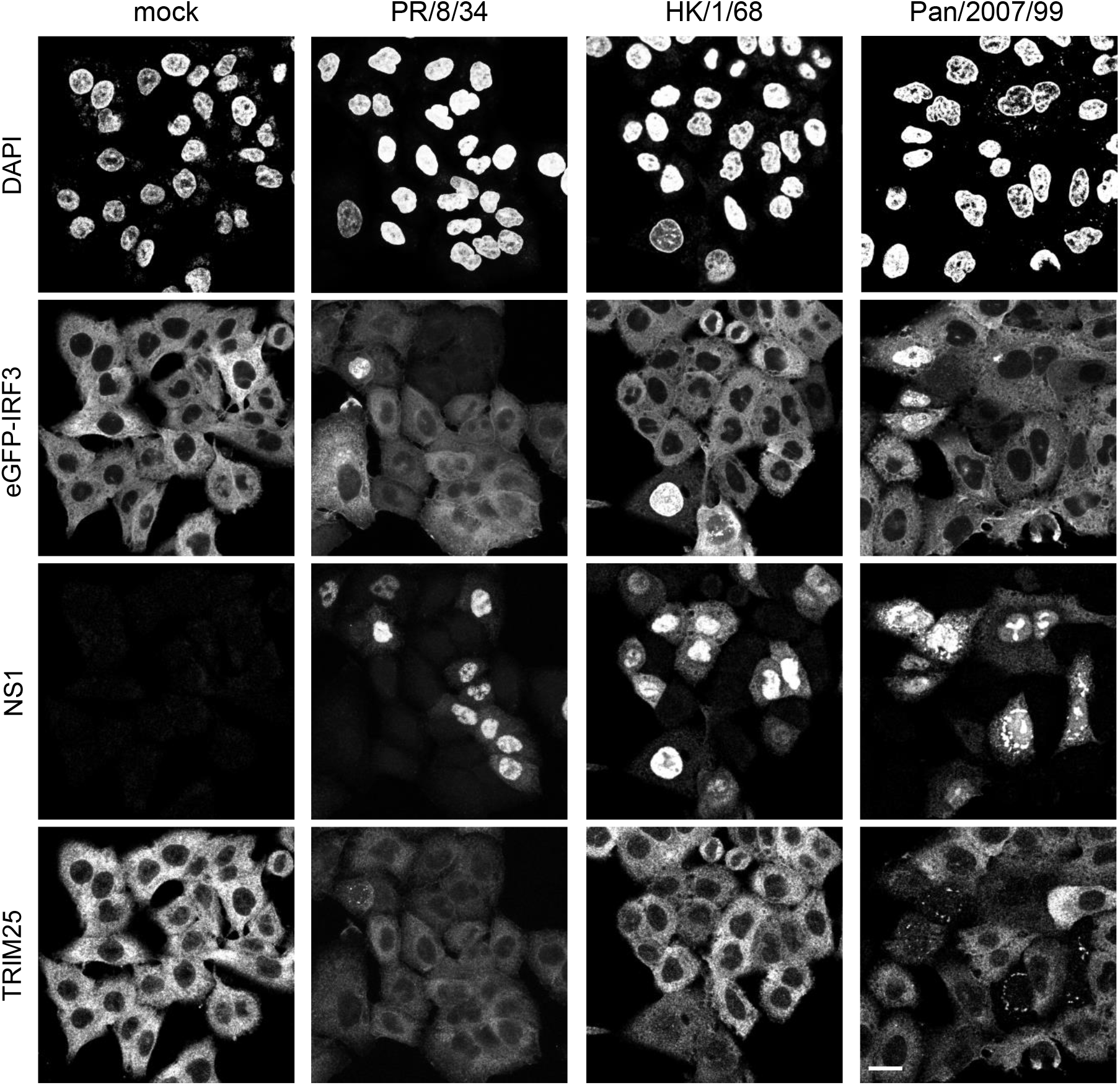
IAV infected A549-eGFP-IRF3 cells with IRF-3 activation by nuclear translocation show minor NS1 aggregates compared to non-activated cells at 16 hours post infection. A549-eGFP-IRF3 cells subjected for immunofluorescence were mock, PR/8/34, HK/1/68, and Pan/2007/99 infected for 16 hours (MOI = 1). Immunostaining was performed using the following first and secondary antibody combinations: mouse anti-NS1/goat anti-mouse Alexa568 and rabbit anti-TRIM25/goat anti-rabbit Alexa647. Image acquisition was performed using an inverted confocal microscope. Images are shown as single planes with a pixel size of 120 nm. Scale bar corresponds to 20 μm.

**Figure S4.**
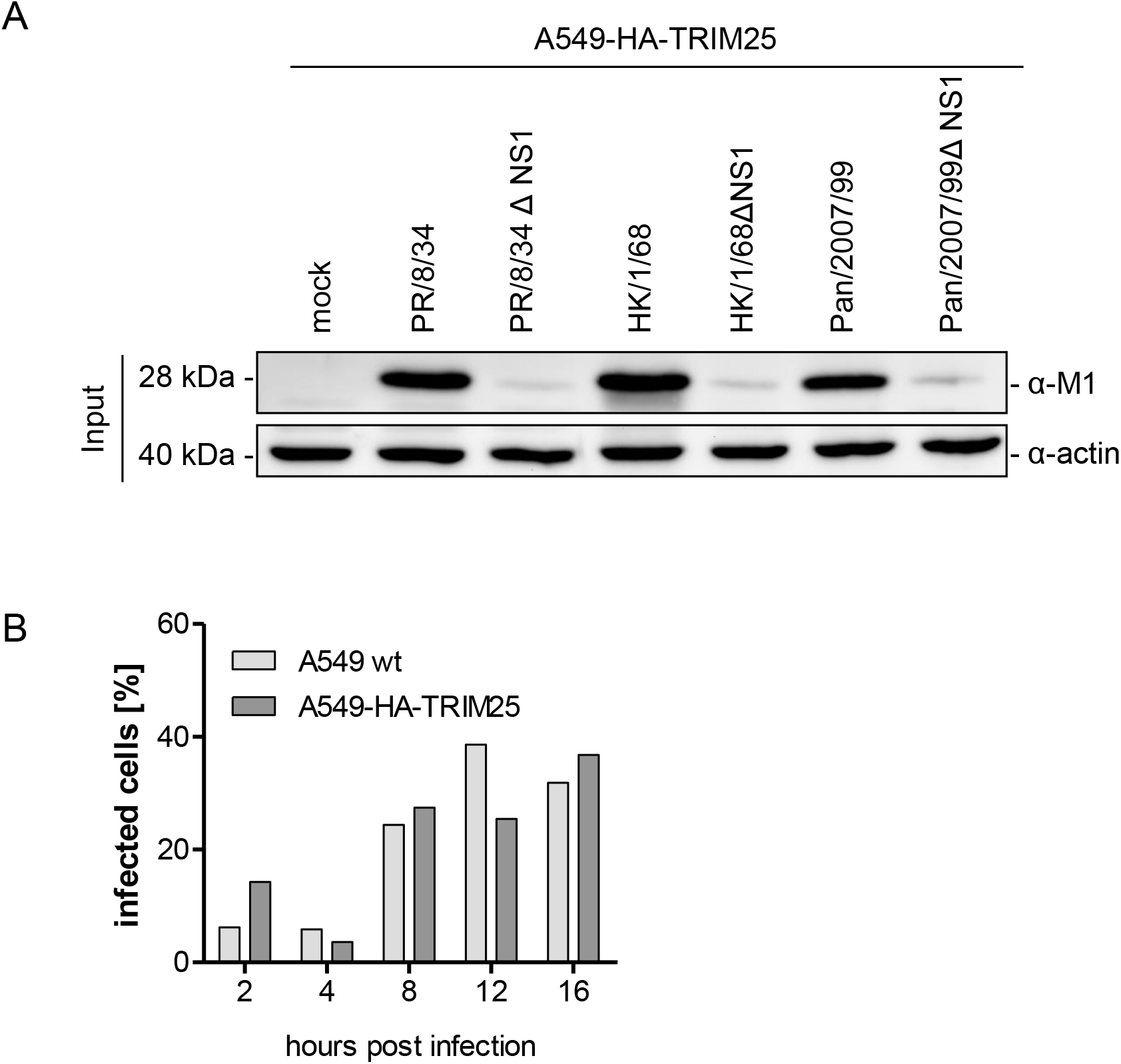
The production of matrix protein 1 is decreased in IAV □NS1 variants compared to corresponding wt strains in A549-HA-TRIM25 cells. **(A)** A549-HA-TRIM25 cells were infected with PR/8/34, HK/1/68, and Pan/2007/99 and their respective ΔNS1 variants. Whole cell lysates were prepared 16 hours post infection. Mock infected control shows no M1 signals. Actin was used as loading control. Panels for the input are derived from a single cropped blot with lanes 1-8 belonging to the same membrane and from the same experiment, respectively. **(B)** A549 wt and A549-HA-TRIM25 cells subjected for immunofluorescence were mock and Pan/2007/99 infected for indicated times (MOI = 1). Fluorescence images were acquired using a wide-field microscope. Infection rate was calculated for each cell line by determining the total cell number based on nucleus counterstaining and the proportion of infected cells being positive for nuclear NP applying a semiautomated analysis approach, (n: A549 wt, 2 hrs = 129 cells, 4 hrs = 153 cells, 8 hrs = 509 cells, 12 hrs = 114 cells, 16 hrs = 436 cells; A549-HA-TRIM25, 2 hrs = 147 cells, 4 hrs = 193 cells, 8 hrs = 175 cells, 12 hrs = 232 cells, 16 hrs = 125 cells)

**Figure S5.**
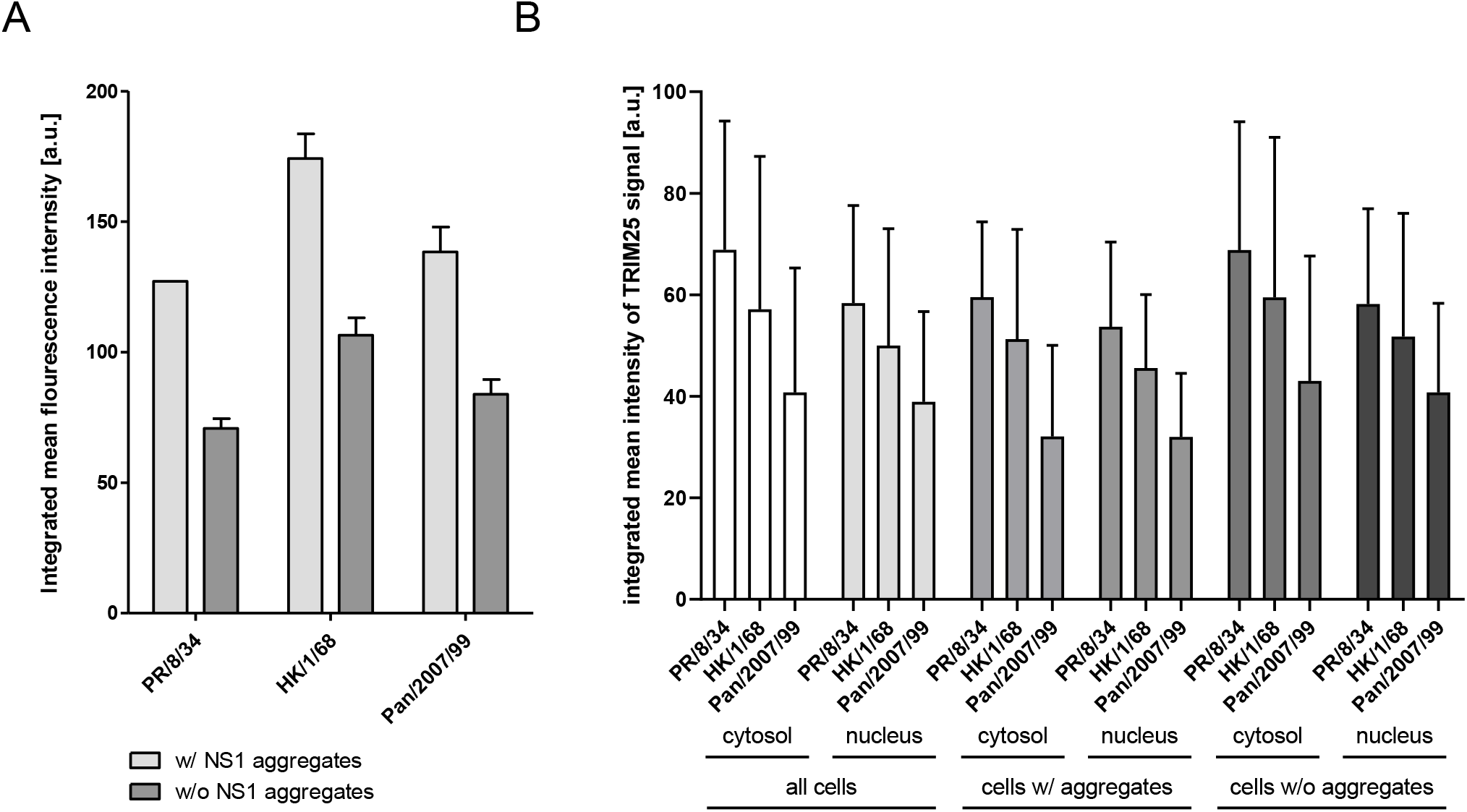
Integrated mean fluorescence intensity of IAV non-structural protein 1 differs between cells with and without cytosolic NS1 aggregates while TRIM25 signals show no significant difference. Quantitative analysis is based on maximal projections of 500 nm z-stacks of PR/8/34, HK/1/68, and Pan/2007/99 infected A549-eGFP-IRF3 cells 16 hours post infection (MOI = 1). **(A)** Graph shows the cytosolic integrated fluorescence intensity of NS1 in cells with and without cytosolic NS1 aggregates in PR/8/34, HK/1/68, and Pan/2007/99 infected A549-eGFP-IRF3 cells. The abundance of NS1 aggregates was analyzed based on NS1 signals in the extra-nuclear space. Results represent the mean (SD) of three independent experiments (n: PR/8/34 = 675 cells, HK/1/68 = 499 cells, Pan/2007/99 = 474 cells). **(B)** Graph shows the integrated fluorescence intensity of TRIM25 in cells with and without cytosolic NS1 aggregates in PR/8/34, HK/1/68, and Pan/2007/99 infected A549-eGFP-IRF3 cells for the cytosol and nucleus, respectively.

